# Endocytosis shapes extracellular chemical gradients in autonomous cell–cell attraction

**DOI:** 10.64898/2026.03.31.715676

**Authors:** Jeremy Barrios, Andrew Goetz, Susan Legette, Purushottam D. Dixit

## Abstract

Receptor-mediated ligand endocytosis is traditionally viewed as a negative feedback mechanism for signal attenuation. Here we show that ligand removal can paradoxically enhance directional information in autonomous cell–cell attraction. Many cell systems migrate toward one another in the absence of externally imposed gradients, implying that secretion, diffusion, and uptake must themselves generate usable directional cues. We develop a surface-resolved theory of a finite-sized detector exposed to a nearby source and derive analytical expressions for the steady-state ligand field. The resulting concentration profiles are governed by a single dimensionless Damköhler number that compares receptor-mediated endocytosis to diffusive ligand transport. Increasing ligand removal lowers extracellular ligand concentrations and reduces absolute concentration differences across the detector surface, but preferentially enhances relative surface anisotropy. Thus, destroying the signal can increase the usable information encoded in relative gradients. Incorporating nonlinear downstream processing reveals a tradeoff between contrast enhancement and signal depletion that yields a well-defined optimal endocytosis rate, in a regime consistent with experimentally measured receptor internalization kinetics. These results recast receptor-mediated endocytosis as an extracellular information-processing mechanism that reshapes self-generated gradients to enhance directional information.

## Introduction

Eukaryotic chemotaxis^1^, the directed movement of eukaryotic cells in response to spatial inhomogeneous chemical cues, underlies essential biological processes such as developmental patterning and immune surveillance^2^, as well as pathological behaviors including cancer invasion and metastasis^3^. A central problem in quantitative cell biology has therefore been to understand how cells reliably extract directional information from chemical signals, particularly when gradients are shallow and noisy^1^.

A dominant theme in chemotaxis research is that cells migrate in stable guidance cues organized at the tissue level. Early theoretical work by Crick^4^ showed that localized ligand production combined with distributed degradation can generate persistent concentration profiles over characteristic diffusion lengths, and subsequent extensions established how feedback^5^, source–sink organization^6^, and tissue geometry^7^ shape gradient profiles over long distances. Together, these studies have motivated a vast body of experimental and theoretical studies focused on how cells sense and respond to fixed, externally imposed chemical fields^8–14^.

Notably, an increasing number of experiments over the past two decades show that cells can cluster together and migrate toward one another even in the absence any external gradient. In developing pancreatic tissue, precursor cell clusters secrete fibroblast growth factor ligands that promote their own aggregation into hormone-expressing clusters^15^. Epithelial cells induced into migratory states extend persistent protrusions toward distant neighbors and assemble into multicellular aggregates, which themselves migrate toward other aggregates over tens to hundreds of microns^16,17^. Similarly, invasive cancer cell lines exhibit robust cluster–cluster attraction in the absence of externally imposed spatial chemical cues^18^. While the molecular mechanisms underlying these behaviors are not always clear, their actionat-a-distance character strongly suggests communication mediated by secreted diffusible ligands. Taken together, these systems pose a basic physical question: how do cells generate surface-resolved directional information at cell–cell length scales when no external guidance cue is provided?

Receptor-mediated endocytosis and subsequent degradation of ligands is known to play an essential role in shaping externally imposed gradients sensed by migrating cells, primarily by acting as a clearance mechanism that sets effective length scales and stabilizes tissue-scale profiles^4–7^. Similarly, endocytosis is also key in interpretation of gradients^12^; blocking endocytosis disrupts chemotaxis^19–22^. By contrast, its role in sculpting the gradient on the cell surface in autonomous cell–cell communication has not been explored. Most theoretical descriptions of autonomous cell–cell communication adopt phenomenological frameworks in which cells are treated as point-like sources and detectors, and the ligand concentration is evaluated only in the bulk, without resolving spatial variation along the cell surface^23–25^. Consequently, it remains unclear how biophysical and signaling parameters, such as ligand secretion rates, ligand diffusivity, receptor density and affinity, and receptor-mediated endocytosis kinetics, jointly determine the surface-resolved directional information available to a cell, or under what conditions such information is sufficient to drive robust attraction.

To address this gap, we develop a minimal, surface-resolved theory of autonomous gradient generation based solely on secretion, diffusion, and receptor-mediated endocytosis. Using an idealized geometry, we derive analytical expressions for the ligand concentration in the extracellular medium and along the surface of a finite-sized detector. This analysis reveals that the entire family of self-generated concentration profiles is governed by a single dimensionless control parameter, a Damköhler number that integrates ligand transport with kinetics of receptor-mediated endocytosis.

Our analysis reveals a fundamental tradeoff in autonomous gradient generation. Increasing receptormediated endocytosis reduces overall ligand abundance and therefore weakens absolute concentration differences across the detector. Paradoxically, the same process sharpens relative ligand anisotropy along the cell surface, enhancing directional contrast in relative gradients, the quantity effectively sensed by many chemotactic circuits^1,11,12^. Because relative sensing typically operates only over a finite dynamic range^12,26^, endocytosis both amplifies directional information and regulates whether the detector operates within this regime. Incorporating a phenomenological model of downstream signal processing predicts a well-defined optimal endocytosis rate, which lies close to experimentally measured rates of growthfactor receptors and G protein-coupled receptors^27,28^.

Together, these results establish a minimal physical framework for autonomous cell–cell attraction at a distance. Our work reframes gradient formation as an extracellular information-processing problem, in which surface-mediated ligand removal tunes the balance between signal amplitude and contrast. This perspective provides a mechanistic foundation for understanding how cells can generate and sense guidance cues without relying on externally imposed gradients.

## Results

We begin by formulating a minimal physical model for autonomous gradient generation in an idealized geometry (Fig. 1). A finite-sized detector of radius *A*, representing a single cell or a compact cell cluster, is placed at the origin. A second cell or cluster, treated as a point source, is located a distance *R* away along a fixed axis and secretes a diffusible ligand at a constant rate. The ligand diffuses in the extracellular medium with diffusion constant *D* and is removed at the detector surface via receptor-mediated endocytosis. The detector may also secrete ligand uniformly from its surface. Owing to the axial symmetry of this configuration, the steady-state ligand concentration depends only on the radial distance from the detector and the polar angle relative to the source.

**FIG. 1.**
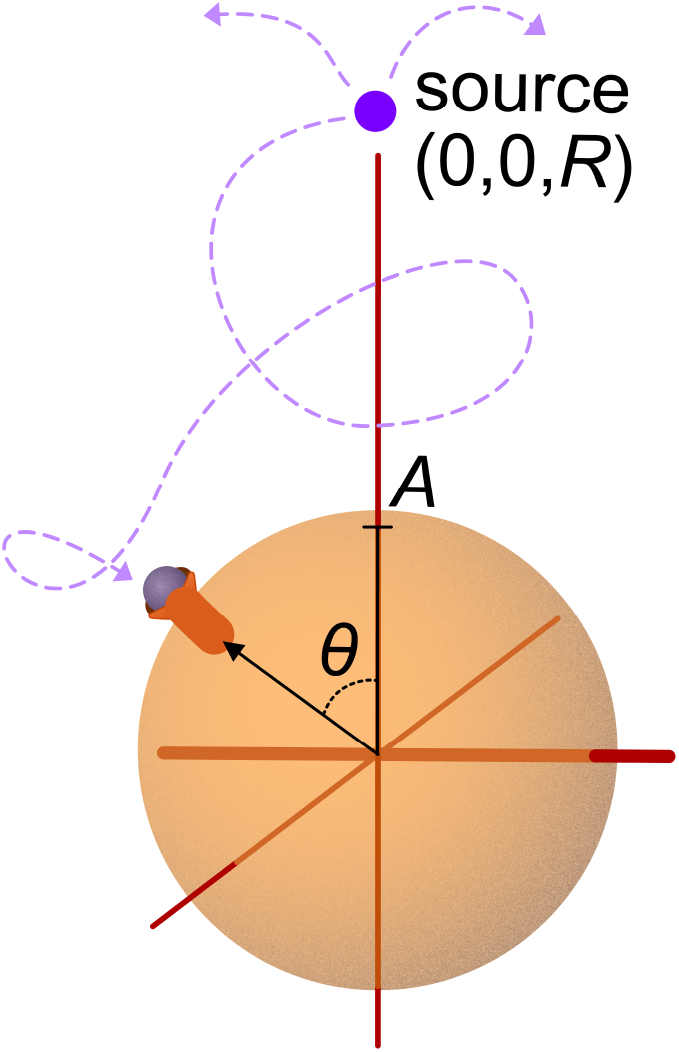
Schematic of the reaction-diffusion model. A spherical detector of radius *A* is situated at origin. A point source of the chemoattractant is situated at (0, 0, *R*). The ligand diffuses with a diffusion constant *D* and is taken up and degraded by the detector at a rate *k*_*c*_. Ligands (pink circles) diffuse from the source and are degradation via receptormediated endocytosis on the surfaces of the detector.

At steady state, the extracellular ligand concentration satisfies the diffusion equation with a localized source,

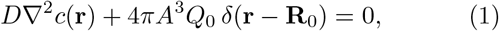

where the point source at position **R**_0_ represents a cell or cell cluster that secretes ligand at a constant rate. Here *Q*_0_ is the source term has units of concentration per unit time and sets the strength of ligand secretion by the emitter. Far from the cells, the ligand concentration vanishes, reflecting dilution into the surrounding medium.

At the surface of the detector, ligand dynamics are governed by a Robin boundary condition that balances diffusive flux with receptor-mediated endocytosis and possible surface secretion,

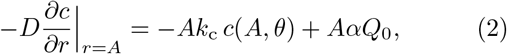

where *k*_c_ is the effective rate of ligand removal via receptor-mediated endocytosis and *αQ*_0_ denotes uniform ligand secretion from the detector surface. This boundary condition captures the essential physics of autonomous gradient generation: ligand removal occurs locally at the detector boundary, while secretion and diffusion act in the bulk.

Before solving Eqs. (1)–(2), it is useful to rewrite the problem in dimensionless form to identify the minimal set of parameters controlling gradient formation. We measure distances in units of the detector radius, *λ* = *r/A*, and define the dimensionless source separation *ρ* = *R/A*. Writing the ligand concentration as *χ*(*λ, θ*) = *c*(*r, θ*)*/ζ*, with

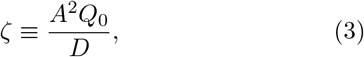

rescales concentrations by the characteristic level set by secretion and diffusion.
In these variables, the governing equations depend on a single dimensionless control parameter,

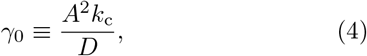

a Damköhler number that compares the rate of receptormediated endocytosis at the detector surface to diffusive transport in the extracellular medium. The relative secretion rate of the detector is captured by the dimensionless ratio *α* = *Q*_1_*/Q*_0_. Expressed in this form, the steadystate ligand profile is fully specified by *ρ, α*, and *γ*_0_, revealing that autonomous gradient generation is governed by a small number of physically transparent parameters. The full analytical solution to Eqs. (1)–(2) is given in the Supplementary Materials. Here we briefly summarize its structure. Owing to axial symmetry about the source–detector axis, the steady-state ligand concentration depends only on the radial distance from the detector and the polar angle relative to the source direction. The solution naturally separates into two regions: an inner region extending from the detector surface to the source location, and an outer region extending from the source to infinity. In each region, the concentration can be expanded in Legendre polynomials,

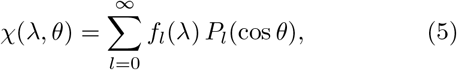

with radial mode functions *f*_*l*_(*λ*) determined by continuity of the concentration, the discontinuity in flux at the source, and the Robin boundary condition at the detector surface. This representation makes explicit how different angular modes contribute to surface anisotropy and provides a direct route to quantifying directional information sensed by the detector.

### Parameter ranges

Before examining the predictions of the model, we identify biologically relevant ranges for the parameters that control ligand transport and sensing. The typical linear size of a eukaryotic cell lies in the range *A ∼*10–60 *µ*m^29^. Diffusible ligands such as growth factors have diffusion constants of order *D ∼*50 *µ*m^2^ s^−129^. Ligand secretion rates are estimated to be in the range 10^3^–10^4^ molecules per cell per minute^30^. The typical separation between communicating cells or compact cell clusters is on the order of a few to ten cell diameters, *R∼* 5–10 *A*^17,18^. The size of such clusters is typically *N*_cell_ *∼*10–100 cells^17,18^.

When non-transformed epithelial cells acquire motility through epithelial to mesenchymal transition (EMT), ligand secretion is often down-regulated^31^. We therefore focus on relative detector secretion rates *α ≤* 1. Unless stated otherwise, we set *α* = 1, corresponding to symmetric secretion by the source and detector.

The key dimensionless control parameter in the model is the Damköhler number *γ*_0_ = *A*^2^*k*_c_*/D*, which compares the rate of receptor-mediated ligand removal at the detector surface to diffusive transport in the extracellular medium. As shown in the Supplementary Materials, *γ*_0_ can be expressed in terms of microscopic signaling parameters as

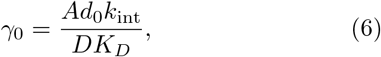

where *d*_0_*∼* 100 *µ*m^−2^ is the surface density of receptors, *K*_*D*_*∼* 0.1–1 nM is the equilibrium dissociation constant of the ligand–receptor interaction, and *k*_int_ *∼* 10^−3^– 10^−2^ s^−1^ is the internalization rate of ligand-bound receptors^27–29,32^. Together, these values suggest that *γ*_0_ spans a broad biologically accessible range, *γ*_0_ *∼*10^−3^–10^1^.

Finally, combining the above estimates yields characteristic concentration scales *ζ* = *A*^2^*Q*_0_*/D* in the range *ζ ∼*10^−3^–1 nM. In the following, unless stated otherwise, we focus on the representative parameter set *ρ* = 4, *α* = 1, and *γ*_0_ = 1, which lies well within the experimentally relevant regime.

### Concentration profile in space

Figure 2(a) shows a representative two-dimensional concentration profile *χ*(*λ, θ*) obtained from the analytical solution. As expected, ligand concentration increases toward the emitting source and decreases far from it. Near the detector, however, the profile exhibits qualitatively different behavior depending on direction. Along the axis pointing away from the source (*θ* = *π*), the concentration decreases monotonically with distance from the detector. In contrast, along the axis pointing toward the source (*θ* = 0), the concentration can be non-monotonic, reflecting the competition between ligand secretion by the detector and endocytotic removal at its surface.

**FIG. 2.**
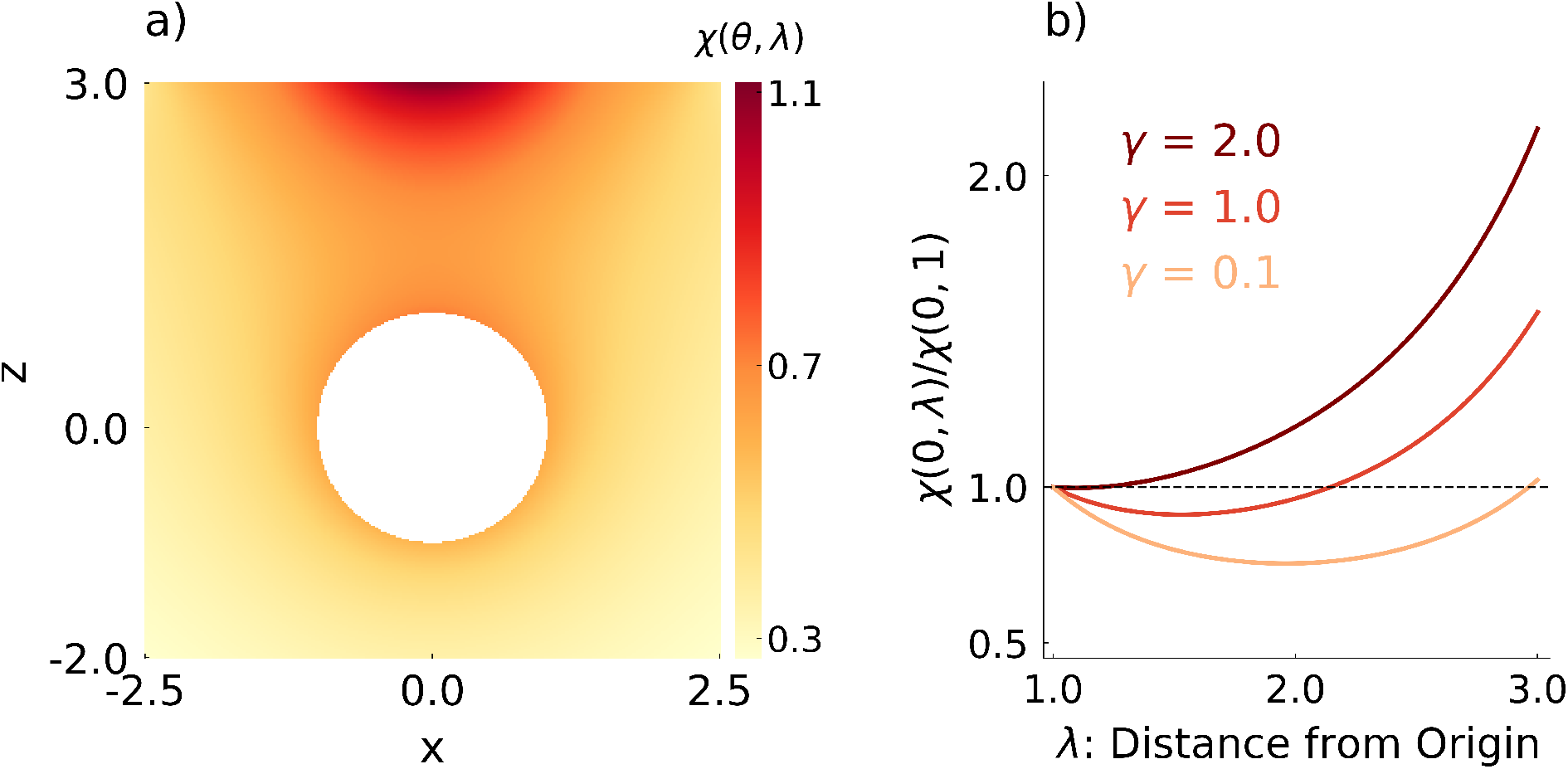
Concentration profiles in space and on the surface of the detector. (a) A two-dimensional dimensionless concentration profile (*χ*(*λ, θ*)) of the ligand. The detector of unit dimensionless radius is situated at the origin. A point source is situated at *ρ* = 4. We have assumed *γ*_0_ = 1 and *α* = 1. (b) Profile of the dimensionless concentration at *θ* = 0 as a function of *λ* for different values of *γ*_0_.

This competition is controlled by the Damköhler number *γ*_0_. As shown in Fig. 2(b), increasing receptormediated endocytosis suppresses the outward flux due to detector secretion, eventually eliminating the nonmonotonicity and producing a concentration profile that increases monotonically away from the detector. These spatial profiles illustrate how surface-localized ligand removal reshapes the extracellular field in a directiondependent manner, even in the absence of any externally imposed gradient.

### Concentration anisotropy on the cell surface

To quantify directional information available to the detector, we focus on the anisotropy of ligand concentration along its surface. Specifically, we define the mean surface concentration on the hemisphere facing the source (0 *≤ θ ≤ π/*2),

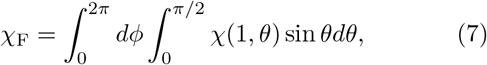

and the corresponding mean concentration on the opposite hemisphere (*π/*2 *≤ θ ≤ π*),

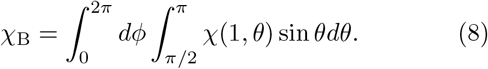

These quantities provide coarse-grained measures of the ligand signal at the front and back of the detector and are natural observables for downstream chemotactic circuits that integrate receptor occupancy over finite regions of the cell surface.

A natural measure of directional signal strength is the absolute difference in ligand concentration across the detector surface,

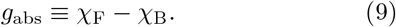

Figure 3(a) shows *g*_abs_ as a function of *γ*_0_ for several source–detector separations. In all cases, the absolute gradient decreases monotonically with increasing receptor-mediated endocytosis. This behavior is intuitive: stronger endocytosis lowers ligand abundance throughout the extracellular medium, reducing concentration differences across the detector surface even as it sharpens spatial structure. As a result, when gradients are quantified purely in absolute terms, increased ligand removal appears detrimental to directional sensing.

**FIG. 3.**
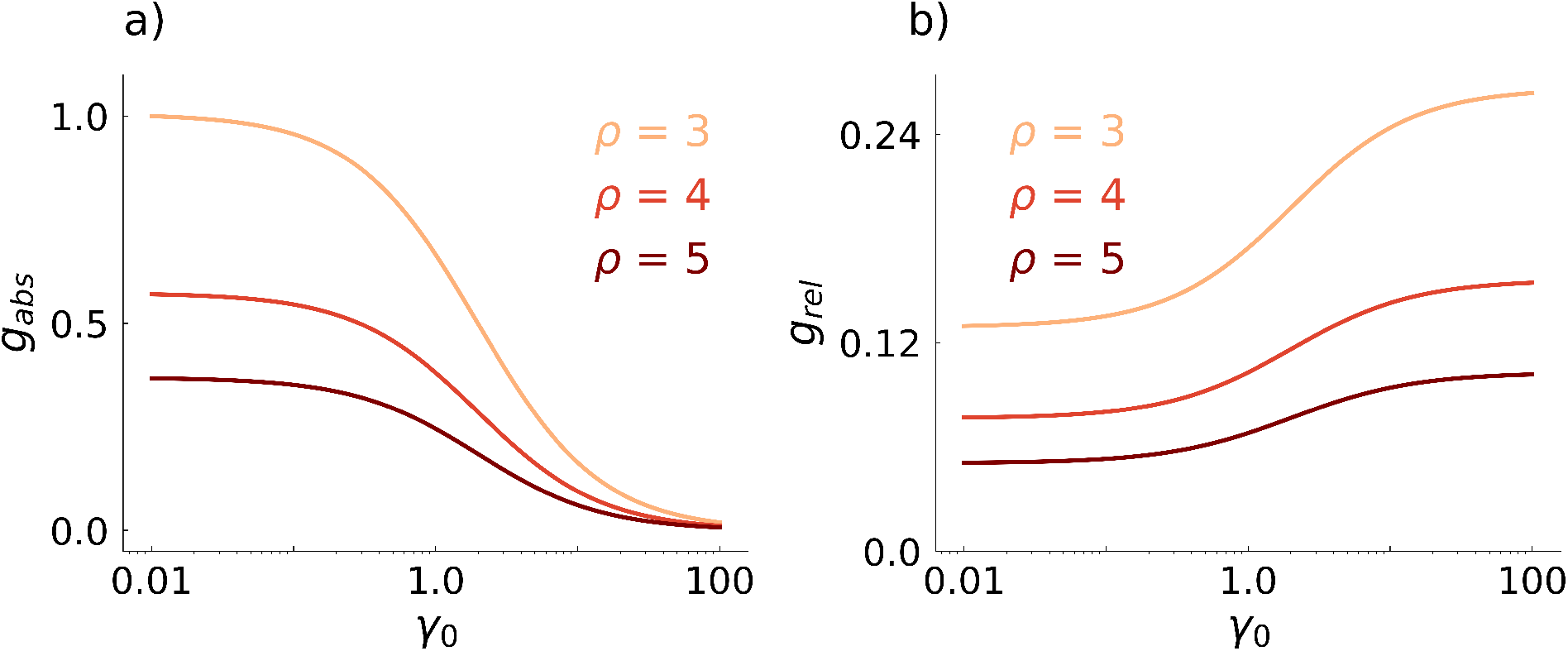
Absolute and relative ligand gradients as a function of ligand degradation. (a) The absolute gradient *g*_abs_ as a function of *γ*_0_ for different values of *ρ*. (b) The relative gradient *g*_rel_ as a function of *γ*_0_ for different values of *ρ*.

While absolute concentration differences decrease with increasing receptor-mediated endocytosis, many chemotactic systems respond not to absolute signals but to relative differences across the cell surface^10,12,33,34^. We therefore define the relative gradient as

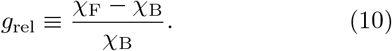

Figure 3(b) shows *g*_rel_ as a function of *γ*_0_ for several source–detector separations. In contrast to the absolute gradient, the relative gradient increases monotonically with increasing endocytosis over a broad parameter range. Compared to the limit of vanishing endocytosis, strong receptor-mediated ligand removal can enhance the relative surface anisotropy by up to a factor of two. Notably, the benefit of endocytosis becomes relevant around *γ*_0_ = 1, well within the ranges predicted by typical signaling parameters (dashed black box).

This behavior highlights a central and counterintuitive feature of autonomous gradient generation: degrading the very signal being detected sharpens the directional contrast available to the detector. Physically, surface-localized ligand removal suppresses the isotropic background concentration more strongly than the anisotropic component induced by the distant source, thereby enhancing relative differences across the detector surface.

### Downstream processing predicts optimal endocytosis rate

If receptor-mediated endocytosis sharpens relative gradients, a natural question is why cells do not simply maximize ligand removal. One reason is that increasing endocytosis, while enhancing relative anisotropy, simultaneously reduces the absolute ligand concentration at the detector surface. Because downstream chemotactic circuits typically respond nonlinearly to ligand exposure, relative sensing is effective only over a finite dynamic range of background signal levels. At very low concentrations the response is negligible, while at high concentrations it saturates.

To capture this constraint in a minimal way, we introduce a phenomenological output that maps surface ligand levels onto a directional response,

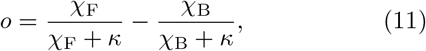

where *κ* sets the midpoint of the downstream activation curve. This form captures the qualitative behavior of many adaptive chemotactic circuits that respond to relative differences only within a limited concentration window (top schematic in Figure 4). Typically, the sensitive ligand concentration range *c*_FCD_ for fold change detection, centered around *κ*, is a couple of orders of magnitude lower than the equilibrium dissociation constant of the ligand (*c*_FCD_*∼* 0.01*K*_*D*_)^12,26^. Since *K*_*D*_*∼* 0.1 − 1nM and *ζ* = 10^−3^− 1nM, This suggests that the typical values of the dimensionless concentration *κ* = *c*_FCD_*/ζ ∼* 10^−3^ − 10^1^. As shown in Fig. 4, the output *o* is maximized at an intermediate value of *γ*_0_, reflecting a balance between contrast enhancement and signal depletion. Notably, the optimal endocytosis rate falls within the experimentally measured range of receptor internalization rates, suggesting that biological systems may operate near regimes that optimize directional information rather than absolute signal strength.

**FIG. 4.**
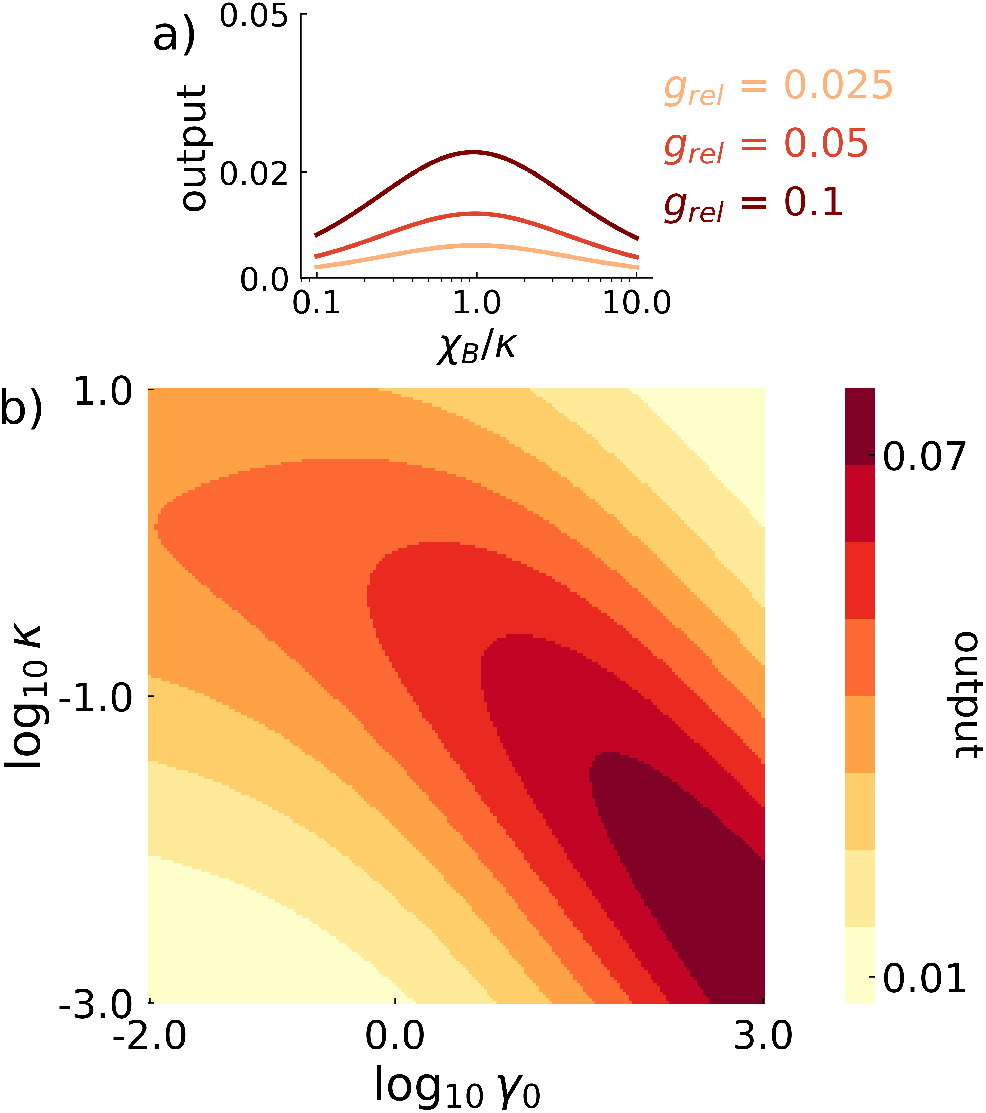
Downstream processing of relative gradients predicts an optimal degradation rate. (top) A schematic of the output as a function of background ligand levels *χ*_*B*_ for increasing values of the relative gradient *g*_rel_. (bottom) Heatmap of the output of the downstream circuit as a function of midpoint of activation *κ* and *γ*_0_.

Together, these results establish a minimal and physically transparent mechanism by which cells can generate and sense directional cues in the absence of any externally imposed gradient. By resolving ligand concentrations along the surface of a finite-sized detector, we show that receptor-mediated endocytosis plays a dual role: it suppresses absolute signal levels while simultaneously enhancing relative surface anisotropy. This tradeoff leads naturally to an optimal endocytosis rate once downstream nonlinearities are taken into account. Importantly, all of these behaviors are controlled by a small number of dimensionless parameters, with the Damköhler number *γ*_0_ acting as the primary control knob. These findings demonstrate that secretion, diffusion, and receptor-mediated endocytosis alone are sufficient to generate robust, surface-resolved directional information at cell–cell length scales.

## Discussion

Receptor-mediated endocytosis is traditionally viewed as a mechanism for attenuating or clearing extracellular signals. In the context of chemotaxis, it is well known to shape tissue-scale gradients by acting as a sink that stabilizes long-range concentration profiles^4–7^. Here, by resolving ligand concentrations along the surface of a finite-sized detector, we show that endocytosis plays a fundamentally different and previously underappreciated role in autonomous cell–cell communication. Rather than merely suppressing signals, surface-localized ligand removal actively reshapes the spatial structure of self-generated gradients, suppressing absolute concentrations while sharpening relative surface anisotropy, the quantity most directly relevant for directional sensing^10,12,33,34^.

A central implication of our results is that secretion, diffusion, and receptor-mediated endocytosis alone are sufficient to generate usable directional information at cell–cell length scales, even in the absence of any externally imposed guidance cue. Our work shows that finite size and boundary-mediated ligand removal are sufficient to break symmetry and generate surface-resolved anisotropy, without invoking additional feedback or spatial patterning mechanisms.

The paradoxical enhancement of relative gradients by endocytosis highlights an important distinction between signal amplitude and information content. While increased ligand removal lowers absolute concentrations throughout the extracellular medium, it preferentially suppresses the background relative to the directional component induced by a distant source. In this sense, receptor-mediated endocytosis functions as an information-processing element that tunes the statistics of the signal before it is decoded by downstream pathways. Importantly, relative gradient enhancement does not imply that endocytosis should be maximized. Once downstream nonlinearities are taken into account, an optimal internalization rate emerges from a balance between contrast enhancement and signal depletion. This optimality arises naturally from the finite dynamic range of relative sensing, rather than from energetic constraints alone. Notably, the predicted optimal regime overlaps with experimentally measured internalization rates of growth-factor receptors and G protein–coupled receptors^27,28^, suggesting that biological systems may operate near parameter values that optimize directional information rather than absolute signal strength.

Recent theoretical work has shown that signal degradation can similarly enhance directional sensing in a threedimensional settings where the receiver cell secretes enzymes to degrade the ligand in space^35^. These findings highlight a broader principle whereby ligand removal can enhance usable information in diffusive fields. It will be interesting to systematically compare such threedimensional sensing strategies with the surface-resolved mechanism studied here, where boundary-localized uptake reshapes extracellular gradients. Such a comparison may clarify how dimensionality and geometry jointly constrain optimal strategies for gradient formation and sensing in biological systems. We leave this exploration for future studies.

Our framework makes several experimentally testable predictions. Pharmacological inhibition or genetic perturbation of endocytosis^36^ should increase overall ligand abundance while reducing relative surface anisotropy, leading to weaker directional bias at long intercellular distances despite higher ligand levels. Conversely, moderate enhancement of endocytosis should improve directional sensing over a finite range of separations before signal depletion dominates. Such effects could be tested by measuring migration speed, directional persistence, or clustering dynamics as a function of distance between communicating cells or clusters^17,18^.

An important limitation of the present model is that ligand removal is modeled without explicitly coupling endocytosis to receptor trafficking, recycling, or surface redistribution. In reality, changes in internalization rates can alter receptor density, spatial organization, and signaling competence, which may further modulate gradient sensing^12^. Extending the model to explicitly incorporate receptor dynamics would allow one to explore feedback between extracellular gradient shaping and intracellular signaling architecture.

More broadly, our results suggest a shift in how autonomous chemotaxis is conceptualized. Rather than viewing gradient sensing solely as an intracellular decoding problem^8,11^, our work emphasizes the role of the sensing machinery in shaping the signal itself. This study provides a minimal physical foundation for understanding how cells communicate directionally in the absence of externally imposed cues. Such mechanisms are likely to be relevant for collective migration, tissue organization, and pattern formation in complex multicellular environments.

## I. SUPPLEMENTARY MATERIALS

### A. Analytical solution

We start with the dimensionless equations

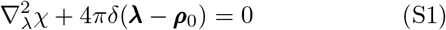

where ***ρ***_0_ = (0, 0, *ρ*) is the dimensionless location of the source. The boundary conditions read:

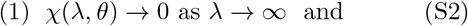

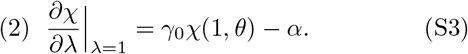

Due to the azimuthal symmetry, the general solution for the problem is given by:

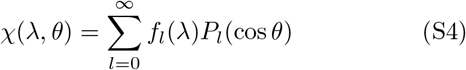

where *P*_*l*_(cos *θ*) are Legendre polynomials and radial functions *f*_*l*_(*λ*) satisfy

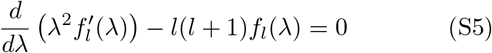

The solutions to Eq. S5 are given by

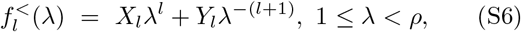

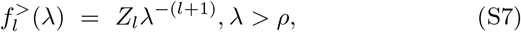

The far field solutions (*λ > ρ*) do not contain *λ*^*l*^ terms to ensure finite concentration as *λ→ ∞* . To ensure continuity of the solution at *λ* = *ρ*, we require:

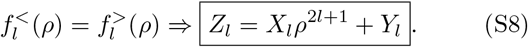

To determine *X*_*l*_ and *Y*_*l*_, we use (1) the jump discontinuity in the radial derivative arising from the points source and (2) the Robin type boundary condition on the surface of the sink. First, we focus on the discontinuity. To evaluate the discontinuity, we start with the governing equation (Eq. **??**):

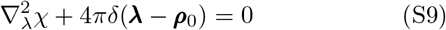

and recognize 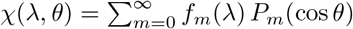. Multiplying both sides by *P*_*l*_(cos *θ*) and integrating over the full solid angle *d*Ω = sin *θdθdϕ*:

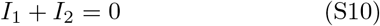

where

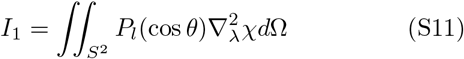

and

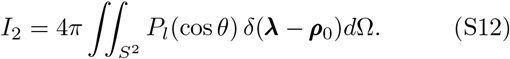

Using

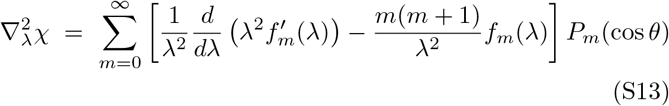

and

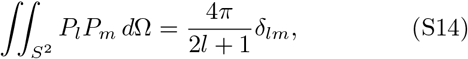

we obtain

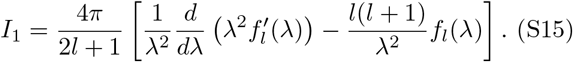

To evaluate *I*_2_, we recognize 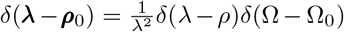, where Ω_0_ is the source direction (*θ*_0_ = 0, *ϕ*_0_ = 0).

Then,

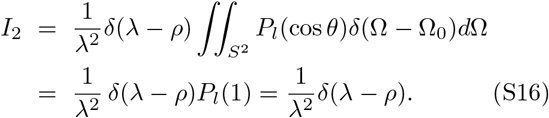

Therefore,

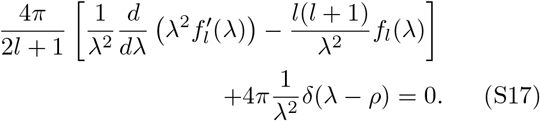

Multiplying by *λ*^2^ and integrating over *λ ∈* (*ρ−ε, ρ* + *ε*):

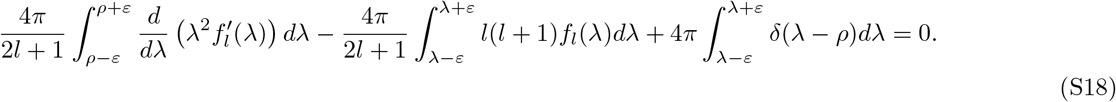

The second term evaluates to zero as *ε →* 0. We have

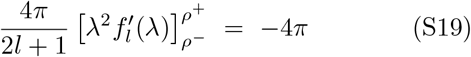

Using Eq. S19, we determine *X*_*l*_. We have

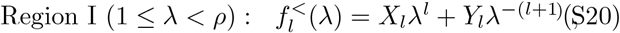

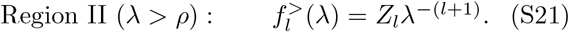

Therefore,

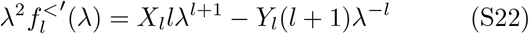

and

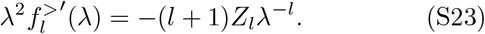

Continuity at *λ* = *ρ* gives *Z*_*l*_ = *X*_*l*_*ρ*^2*l*+1^ + *Y*_*l*_. Therefore,

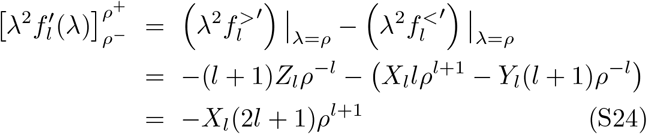

where the *Y*_*l*_ terms cancel using *Z*_*l*_ = *X*_*l*_*ρ*^2*l*+1^+*Y*_*l*_. Therefore, we have

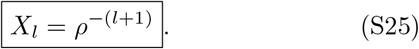

Next, we determine *Y*_*l*_ using the Robin boundary conditions. At *r* = *A*, we impose

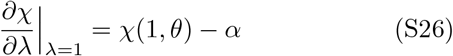

We evaluate the derivatives in region 1,

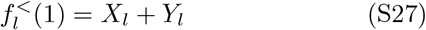

and

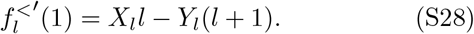

Next, we project the boundary condition onto *P*_*l*_(cos *θ*). Since *− αγ*_0_ does not depend on *θ*, it contributes only to the *l* = 0 mode. Therefore for *l >* 0:

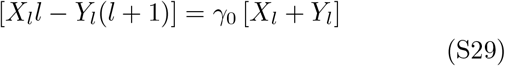

Solving for *Y*_*l*_ yields

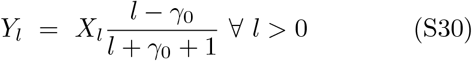

For *l* = 0,

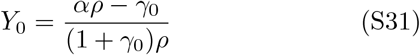

*Z*_*l*_ can be obtained from *X*_*l*_ and *Y*_*l*_ using Eq. S8.

### B. Expressing *γ*_0_ using signaling parameters

Let us consider the boundary condition:

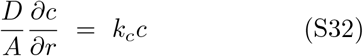

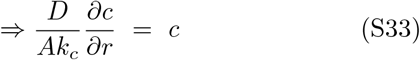

The total flux of ligands at the surface is (from the environment to the cell)

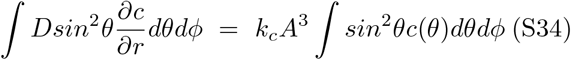

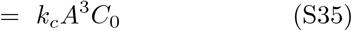

Where

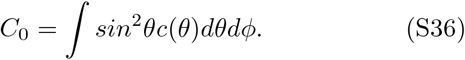

Let us assume that the receptor density is *d*_0_. When the ligand concentration is small, the total number of bound receptors in a small region of the surface is given by 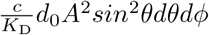 where *K*_D_ is the equilibrium dissociation constant of the ligand. Therefore, we have an expression for the net number of receptors degraded along the cell surface:

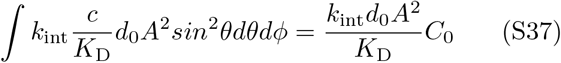

where *k*_int_ is the internalization rate of receptors. Equating the two fluxes, we have

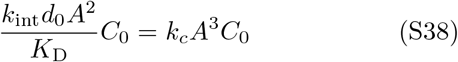

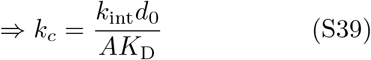

The relevant dimensionless constant is *γ*_0_ = *A*^2^*k*_*c*_*/K*_*D*_, given by

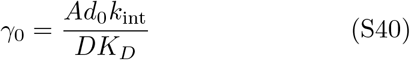

